# MRI-guided histology of TDP-43 knock-in mice implicates parvalbumin interneuron loss, impaired neurogenesis and aberrant neurodevelopment in ALS-FTD

**DOI:** 10.1101/2020.05.24.107177

**Authors:** Ziqiang Lin, Eugene Kim, Mohi Ahmed, Gang Han, Camilla Simmons, Yushi Redhead, Jack Bartlett, Luis Emiliano Pena Altamira, Isobel Callaghan, Matthew A. White, Nisha Singh, Stephen Sawiak, Tara Spires-Jones, Anthony C. Vernon, Michael P. Coleman, Jeremy Green, Christopher Henstridge, Jeff S. Davies, Diana Cash, Jemeen Sreedharan

**Affiliations:** Department of Basic and Clinical Neuroscience, The Maurice Wohl Clinical Neuroscience Institute, Institute of Psychiatry, Psychology and Neuroscience (IoPPN), King’s College London, London, UK; BRAIN Centre (Biomarker Research And Imaging for Neuroscience), Department of Neuroimaging, IoPPN, King’s College London, London, UK; Centre for Craniofacial and Regenerative Biology, Floor 27 Tower Wing, Guy’s Hospital, King’s College London, London, UK; Molecular Neurobiology, Institute of Life Sciences, School of Medicine, Swansea University, Swansea, UK; Centre for Discovery Brain Sciences, University of Edinburgh, Edinburgh, UK; Division of Systems Medicine, School of Medicine, University of Dundee, Dundee, UK; West China School of Medicine, West China Hospital, Sichuan University, Chengdu, China; School of Biomedical Engineering & Imaging Sciences, St Thomas’ Hospital, King’s College London, 4th floor Lambeth Wing, London; Department of Clinical Neurosciences, Cambridge University, Cambridge, UK; Brain Repair Centre, Cambridge University, Cambridge, UK

**Keywords:** frontotemporal dementia, amyotrophic lateral sclerosis, motor neuron disease, TDP-43, parvalbumin, neurogenesis, microglia, magnetic resonance imaging

## Abstract

Amyotrophic lateral sclerosis (ALS) and frontotemporal dementia (FTD) are overlapping neurodegenerative diseases that are increasingly understood to have long prodromal periods. Investigation of these early stages promises to yield valuable biomarkers of disease and will be key to understanding mechanisms underlying the genesis of ALS-FTD. Here, we use *in vivo* magnetic resonance imaging (MRI), histology and computed tomography to identify structural and cellular readouts of early stage disease in the TDP-43^Q331K^ knock-in mouse model of ALS-FTD. Adult mutant mice demonstrated parenchymal volume reductions affecting the frontal lobe and entorhinal cortex in a manner reminiscent of ALS-FTD. Subcortical, cerebellar and brain stem regions were also affected in line with observations in presymptomatic carriers of mutations in *C9orf72*, the commonest genetic cause of both ALS and FTD. Volume loss, as measured by MRI, was also observed in the dentate gyrus (DG) of the hippocampus, along with ventricular enlargement. Guided by these imaging findings, detailed *post-mortem* brain tissue analysis revealed reduced parvalbumin-positive (PV+) interneurons as a potential cellular correlate of MRI changes in mutant mice. By contrast, microglia were in a disease activated state even in the absence of brain volume loss. A reduction in immature neurons was found in the DG, indicative of impaired adult neurogenesis, while a paucity of PV+ interneurons in juvenile mutant mice (P14) suggests that TDP-43^Q331K^ disrupts neurodevelopment. Computerised tomography imaging also showed altered skull morphology in mutants, further suggesting a role for TDP-43^Q331K^ in development. Finally, analysis of human post-mortem prefrontal cortices confirmed a paucity of PV+ interneurons in the prefrontal cortex in cases with both sporadic ALS and ALS linked to *C9orf72* mutations. This study suggests an important role for PV+ interneurons in regional brain vulnerability associated with ALS-FTD, and identifies novel MRI and histological biomarkers that will be of value in assessing the efficacy of putative therapeutics in TDP-43^Q331K^ knock-in mice.

## Introduction

Amyotrophic lateral sclerosis (ALS) and frontotemporal dementia (FTD) exist on a clinicopathological and genetic disease spectrum (ALS-FTD). The loss of upper and lower motor neurons in ALS leads to paralysis, while FTD is characterized by frontal and temporal lobar degeneration causing executive dysfunction, language impairment and behavioural changes (Ahmed et al., 2016). From the point at which patients with ALS-FTD describe their first symptoms the decline is often rapid, with death ensuing within 3-5 years. However, it is increasingly apparent that patients can have seemingly asymptomatic prodromes that can extend for decades prior to clinical onset. This conclusion is based on observations from recent studies using sensitive magnetic resonance imaging (MRI) methods, which are capable of detecting subtle deviations in brain structure or function in pre-symptomatic carriers of ALS-FTD-linked gene mutations including *C9orf72* (Menke et al., 2017; Bertrand et al., 2018; Cash et al., 2018; Panman et al., 2019). It follows that combining genetics with non-invasive imaging is a powerful approach to identify biomarkers of early disease (Thompson et al., 2010). This is invaluable to enable early diagnosis and early therapeutic intervention to delay onset, slow progression or even prevent disease altogether, thereby providing the greatest possible benefit to patients.

Almost all cases of ALS and half of FTD cases are characterised by pathological processing of the RNA-binding protein TDP-43 (Arai *et al.*, 2006; Neumann *et al.*, 2006; Burrell *et al.*, 2016). Mutations in *TARDBP*, which encodes TDP-43, are also linked with ALS and FTD (Sreedharan *et al.*, 2008; Benajiba *et al.*, 2009). These observations indicate key but undefined roles for TDP-43 in the pathogenesis of ALS-FTD. To elucidate molecular processes underlying the genesis of ALS-FTD it would be ideal to spatially resolve early brain changes *in vivo* and correlate regional atrophy with histological data in TDP-43 preclinical models. Such an approach promises to identify cross-species biomarkers of disease, with the potential to facilitate translational research. *In vivo* brain imaging, for example, offers the potential to directly correlate results from animal disease models with human imaging data. However, few such studies have been conducted in ALS-FTD models of disease, and these have not utilised an unbiased approach to identify affected brain regions of interest for further study.

Ideally, preclinical models should accurately reflect the human genetic state to ensure that observations are not due to artefacts caused by exogenous expression of transgenes. In the case of TDP-43, it is also important to ensure ubiquitous expression including throughout development (Wu *et al.*, 2010). These criteria are fulfilled by the TDP-43^Q331K^ knock-in mouse, which harbours a single human-equivalent point mutation in the endogenous mouse *Tardbp* gene (White *et al.*, 2018). We previously showed that this mutation causes a gain of function due to a loss of TDP-43 autoregulation. We also demonstrated that TDP-43^Q331K^ knock-in mice display progressive behavioural features of FTD indicating dysfunction of frontal and temporal-equivalent brain regions. Here, using *in vivo* MRI, *ex vivo* CT and histology of TDP-43^Q331K^ knock-in mice, we find a striking recapitulation of regional brain atrophy as seen in human ALS-FTD, indicating the value of the TDP-43^Q331K^ knock-in mouse as a translational tool. We then use these MRI results to guide our histological studies, which implicate interneurons, microglia, impaired neurogenesis and aberrant neurodevelopment in early stage disease. Finally, we validate our interneuron observation using histology of human postmortem brain tissues.

## Materials and methods

### Mouse breeding and maintenance

The creation of the TDP-43^Q331K^ knock-in mouse was previously described (White et al., 2018). For the current study, heterozygous (TDP-43^Q331K/+^) mice were intercrossed to generate TDP-43^Q331K/Q331K^, TDP-43^Q331K/+^ and wild-type offspring. All mice were bred on a C57Bl/6J background. Genotyping for the Q331K mutation was performed as described previously (White et al., 2018). All animal experiments were performed under the UK Animal (Scientific Procedures) Act 1986 Amendment Regulations 2012 on Project Licences P023CC39A and P35785FD7. Animals were group-housed in individually ventilated Tecniplast cages within a clean facility. Individual cages contained environmental enrichment items and group sizes of two to five mice were routinely maintained under a 12 h light/dark cycle with ad libitum access to food and water.

### Structural magnetic resonance imaging

*In vivo* magnetic resonance imaging (MRI) was performed on 7-month-old (28-32 weeks, 27.3-38.6 g) male mice (10 TDP-43^Q331K/+^, 10 TDP-43^Q331K/Q331K^ and 15 wild-type littermates) using a 9.4 T horizontal bore Bruker BioSpec 94/20 scanner (Bruker BioSpin, Ettlingen, Germany). An 86-mm volume resonator and a 2×2 phased array surface coil were used for RF transmission and reception respectively. During scanning, the mice were anesthetized with 2-2.5% isoflurane. Their respiration rate and core temperature were monitored and maintained at 60-80 breaths/min and 37±0.5°C (SA Instruments, Inc.). 3D T1-weighted images were acquired using an MPRAGE sequence with the following parameters: echo time (TE) = 3.8 ms, repetition time (TR) = 9.66 ms, inversion time (TI) = 1300 ms, flip angle = 10°, segment TR = 3500 ms, 4 segments, 4 averages, field of view (FOV) = 16×16×8 mm, matrix = 128×128×64, scan time = 40 minutes. One wild-type and one TDP-43^Q331K/+^ mouse were excluded due to poor image quality.

Visual inspection of the MR images revealed that ventricular enlargement is a clear phenotype of homozygous TDP-43^Q331K/Q331K^ mutants. To quantify this using MATLAB (MathWorks, Natick, MA, USA) and FSL (FMRIB, Oxford, UK.), the ventricles of each mouse were semi-automatically segmented through a combination of thresholding, morphological operations, and manual editing if necessary. The volumes of these ventricle masks were compared across genotypes.

Tensor-based morphometry was performed to assess the effects of mutant TDP-43^Q331K^ on local brain volume. First, N4 bias field correction was applied to the MPRAGE images to remove the signal inhomogeneity arising from the surface coil’s nonuniform sensitivity profile. Then, a study-specific template was created using ANTs (antsMultivariateTemplateConstruction2.sh). All subjects were registered to the template via sequential rigid, affine, and nonlinear SyN transforms (antsRegistration). The MPRAGE images and ventricle masks were given equal weight in the registration to improve the normalization of the largely varying ventricle sizes. Two TDP-43^Q331K/Q331K^ mice with frank hydrocephalus were excluded from further analysis because their ventricles could not be satisfactorily normalized to the template.

Jacobian determinant maps of the resultant deformation fields were computed and log-transformed. To compare local volume differences between mutant and wild-type mice, voxelbased nonparametric statistics were performed on the log-transformed Jacobian determinant maps using FSL randomise (5000 permutations, threshold-free cluster enhancement, familywise error (FWE) correction). An ANOVA was performed, followed by pair-wise comparisons between genotypes. We did not correct for global scaling as total brain volumes (i.e. brain plus ventricles) did not differ between genotypes.

The DSURQE mouse brain atlas (The Mouse Imaging Centre, Toronto) (Dorr *et al.*, 2008) was similarly registered to the study-specific template for ROI analysis. For each mouse, ROI volumes were calculated by summing the Jacobian determinant values within each ROI. MATLAB was used to perform two-tailed Mann-Whitney U tests to compare ROI volumes between each pair of groups, controlling for false discovery rate (FDR) using the Benjamini-Hochberg step-up procedure.

### Mouse immunohistochemical studies

#### Tissue preparation

Following MRI, mice were culled by asphyxiation with CO2 followed by cervical dislocation and tissue extraction. The brains were immersion fixed in 4% paraformaldehyde (PFA) at 4°C for 24-48 h, washed in phosphate buffered saline (PBS), and cryoprotected in 30% sucrose in PBS at 4°C for 72 h. The tissues were then embedded in M1 matrix in a silicon mould, frozen on dry ice, sectioned at 35 μm thickness on a cryostat (Leica) and stored in cyroprotectant (25% ethylene glycol, 30% glycerol) at −20°C. Tissue sections were then immunostained as follows (a list of the antibodies used is provided in Supplementary Table 1).

#### Immunostaining

For parvalbumin (PV) immunofluorescent staining cryoprotected sections were washed in PBS for 20 min, 0.2% TritonX-100/PBS for 10 min, and blocked in 5% NGS/0.2% TritonX-100/PBS for 1h at RT. Sections were incubated overnight at 4°C with primary anti-PV antibody diluted in blocking buffer, washed in 0.2% TritonX-100/PBS, then incubated for 2 h at RT with secondary antibody diluted in blocking buffer, then washed in 0.2% TritonX-100/PBS. Sections were subsequently washed in PBS and mounted onto Superfrost Plus slides (VWR, France) with Vectashield HardSet with DAPI (Vector lab).

For Iba1 and Tmem119 co-immunofluorescent staining, antigen retrieval was first performed by heating the samples for 20 min at 80° C in sodium citrate buffer (pH 6.5). Sections were cooled to R/T, washed in distilled water, and blocked and permeabilized in a solution containing 5% BSA, 0.5% Triton X-100 for 1 hour at R/T. Slides were incubated with primary antibody (goat anti-Iba1, Abcam, ab5076, 1:500) for 4° C overnight in twofold-diluted blocking buffer. Secondary antibody (donkey anti-goat, Alexa Fluor 488, 1:500) was applied for 1 hour at R/T. Sections were then washed and blocked again as above and incubated with primary antibody (rabbit anti-Tmem119, Abcam, ab209064, 1:1000) for 4°C overnight. Secondary antibodies were applied for 1 h at R/T (goat anti-rabbit, Alexa Fluor 568, 1:500). Sections were counterstained and mounted with Vectashield HardSet with DAPI (Vector Labs).

For DAB-immunohistochemistry, sections were washed in 0.1M PBS and 0.1M PBS-T. Endogenous peroxidases were quenched by washing in PBS plus 1.5% H2O2 for 20 min. Sections were washed again. Antigen retrieval was performed where needed (i.e. for MBP and Ki67 staining) in sodium citrate at 70°C for 1 h. Sections were blocked in 5% NGS in PBS-T for 1 h, washed, and then incubated overnight at 4°C with primary antibody in PBS-T and 2% NGS solution. Sections were washed again prior to incubation with biotinylated secondary antibody in PBS-T for 70 min. Sections were washed and incubated in ABC solution (Vectastain ABC Elite, PK-6000, Vector Laboratories, USA) for 90 min in the dark, then washed twice more in PBS, and incubated with 0.1M sodium acetate pH 6 for 10 min. Immunoreactivity was developed in DAB (3,3′-Diaminobenzidine) solution followed by two washes in PBS. Sections were mounted and allowed to dry overnight before being dehydrated and de-lipidized in increasing concentrations of ethanol. Sections were incubated in Histoclear (2×3 min; National Diagnostics, USA) and cover-slipped with mounting medium (Sigma Aldrich, UK).

#### Image analyses

For adult mouse immunostaining, images were captured using an Olympus Whole Slide Scanner (VS120, Olympus, Japan) with a x20 objective. Z-stacks of 5 layers were obtained at x20 magnification throughout each section. Images were auto-stitched using Olyvia software (Olympus). Images were analysed using Visiopharm image analysis software (Hoersholm, Denmark) blind to genotype. Anatomically equivalent sections were selected from 4-6 mice of each genotype. ROIs were manually drawn in each chosen section according to the Allen mouse brain atlas (Allen Institute for Brain Science). Each ROI extended through 3-6 sections depending on its size. Each marker was measured using a custom Visiopharm APP via threshold classification and post processing changes. The number of any particular cellular subtype per tissue area (‘cell density’), and the proportion of an area positive for any marker (‘percentage area’), were calculated. The measurements in all sections for each ROI were summed for each mouse (density = total cell number/ total tissue area; percentage area = total marker-positive area/ total tissue area).

For quantification of PV+ interneurons in P14 mouse hippocampi, cells were manually quantified (30μm confocal z-stacks, Leica TCS SP5) from 6 anatomically equivalent sections per animal within a given region using Fiji (ImageJ). Watershed algorithm was used to define the ROI and DAPI+ cells quantified with the parameters – radius:0.5; min level:90-127; max level:127-192; erode/dilate iterations:0 in binary images. Mean pixel intensities were checked for accuracy by redirecting measurements to the original image for overlay with DAPI.

The fold change of immunostaining in all animals for each ROI was calculated relative to the mean value of wild-type controls.

### *Ex vivo* micro-CT imaging and data analysis

For micro-CT, 24-month-old male mice were perfused with 2% paraformaldehyde, and heads were processed as previously described (White *et al.*, 2019). Heads were scanned using a Scanco μCT50 micro-CT scanner (Scanco, Brüttisellen, Switzerland). The specimens were immobilised in 19 mm scanning tubes using cotton gauze and scanned to produce 20 μm voxel size volumes, using X-ray settings of 70kVp, 114uA and a 0.5 mm aluminium filter to attenuate harder X-rays. The scans were automatically scaled at reconstruction using the calibration object provided by the CT manufacturer, consisting of five rods of hydroxyapatite (HA) at concentrations of 0 to 790mg HA/cm3, and the absorption values expressed in Hounsfield Units (HU).

For CT image analysis, 53 landmarks of the skull were obtained by reconstructing a 3D surface of the micro-CT images using Microview software by Parallax Innovations (http://www.parallax-innovations.com/microview.html) followed by genotype-blinded manual placement and orthogonal-view cross-checking. The MorphoJ software package (Hallgrimsson *et al.*, 2007) was used to apply Procrustes superimposition to landmarks for alignment after which shape variation was visualised through principal component analysis. Procrustes Distance Multiple Permutations test (1000 iterations) was used to quantify the significance of shape differences between genotypes. Size differences were compared by measuring the centroid size (square root of the sum of the squared distances from each landmark to the centroid) and significance measured via an unpaired t-test. Videos were generated by taking one representative homozygous mutant and one representative wild-type mouse and using a pipeline as previously described (Toussaint *et al.,* 2019) to morph between them.

### Human tissue studies

#### Patient details

All patients had clinical and electrophysiological evidence of combined upper and lower motor neuron dysfunction and fulfilled the revised El Escorial criteria for a diagnosis of ALS (Brooks et al., 2000). Patients were recruited through the Scottish Motor Neurone Disease Register and data collected in the CARE-MND database (Clinical Audit Research and Evaluation). Ethical approval for this register was obtained from Scotland A Research Ethics Committee 10/MRE00/78 and 15/SS/0216. All clinical data were subsequently extracted from the CARE-MND database.

Patients gave pre-mortem consent for brain and spinal cord donation. All tissue retrievals were in line with the Human Tissue (Scotland) Act, and all procedures have been approved by a national ethics committee. Use of human tissue for post-mortem studies has been reviewed and approved by the Sudden Death Brain Bank ethics committee and the ACCORD medical research ethics committee, AMREC (ACCORD is the Academic and Clinical Central Office for Research and Development, a joint office of the University of Edinburgh and NHS Lothian). Details of all donors used in this study (ages, sex and C9orf72 mutation status) with means for each group can be found in Supplementary tables 2 and 3.

#### Neuropathology

Fresh post-mortem tissue blocks of human brain were fixed in 10% formalin for a minimum of 24 h. Tissue was dehydrated in an ascending alcohol series (70–100%), followed by three xylene washes, all for 4 h each. Next, three paraffin waxing stages (5 h each) were performed to ensure full penetration of the embedding wax and the blocks were cooled. Tissue sections were cut on a Leica microtome at 4 μm and collected on glass slides. All sections were dried at 40°C for at least 24 h before staining. Immunohistochemistry was performed using standard protocols, enhanced using a sensitive avidin/biotin kit (VECTASTAIN ABC Elite) and visualized using DAB as chromogen. The primary antibody was rabbit anti-Parvalbumin (Swant, PV27, 1:1000). Slides were finally counterstained with hematoxylin for 30 s to stain cell nuclei. All staining was performed by investigators blind to clinical diagnosis.

One section per case was analysed and PV+ cell densities were generated using Stereo Investigator. Cortical grey matter was outlined in each section and immune-positive cells identified using an automated colour-based thresholding algorithm. The accuracy of this automated process was assessed on each section by manually checking the identified objects. The number of immuno-positive cells was expressed as a cell density by dividing the total number by the total cortex area, expressed as PV+ cells/mm2. All imaging and analysis were performed by investigators blind to clinical diagnosis.

### Experimental design and statistical analysis

Experimental data were analysed with the operator blind to the genotype of animals. All statistical analyses were conducted, and all figures plotted using Prism 8.3.0 (GraphPad Software, Inc). All statistical comparisons are based on biological replicates unless stated otherwise. All charts show mean ± s.e.m. and statistical tests used are described in the relevant results or figure legends. P-values <0.05 were considered significant for all statistical analyses used unless otherwise indicated.

### Data availability

The authors will make all data available to readers upon reasonable request.

## Results

### Regional parenchymal volume loss and ventricular enlargement on *in vivo* MRI

To identify structural brain changes caused by mutant TDP-43^Q331K^ we performed *in vivo* MRI on 7-month-old male heterozygous (TDP-43^Q331K/+^) and homozygous (TDP-43^Q331K/Q331K^) mutant mice and wild-type littermates. This represents an early symptomatic stage in this model with evidence of executive dysfunction, but no motor neuron loss as previously described (White et al., 2018). MRI revealed that the brain parenchymal volume decreased by 5.0% (Fig. 1A, B). Furthermore, there was a striking 49.7% increase in total ventricular volume in TDP-43^Q331K/Q331K^ mice compared to wild-types (Fig. 1C). There was no significant difference in total brain and ventricular volume combined (Fig. 1D). Furthermore, voxelwise analysis using tensor-based morphometry (TBM) (White et al., 2019) revealed widespread statistically significant (FWE P<0.05) volume reductions in cortical, subcortical and cerebellar regions in TDP-43^Q331K/Q331K^ compared to wild-type mice (Figure 1E).

**Figure 1.**
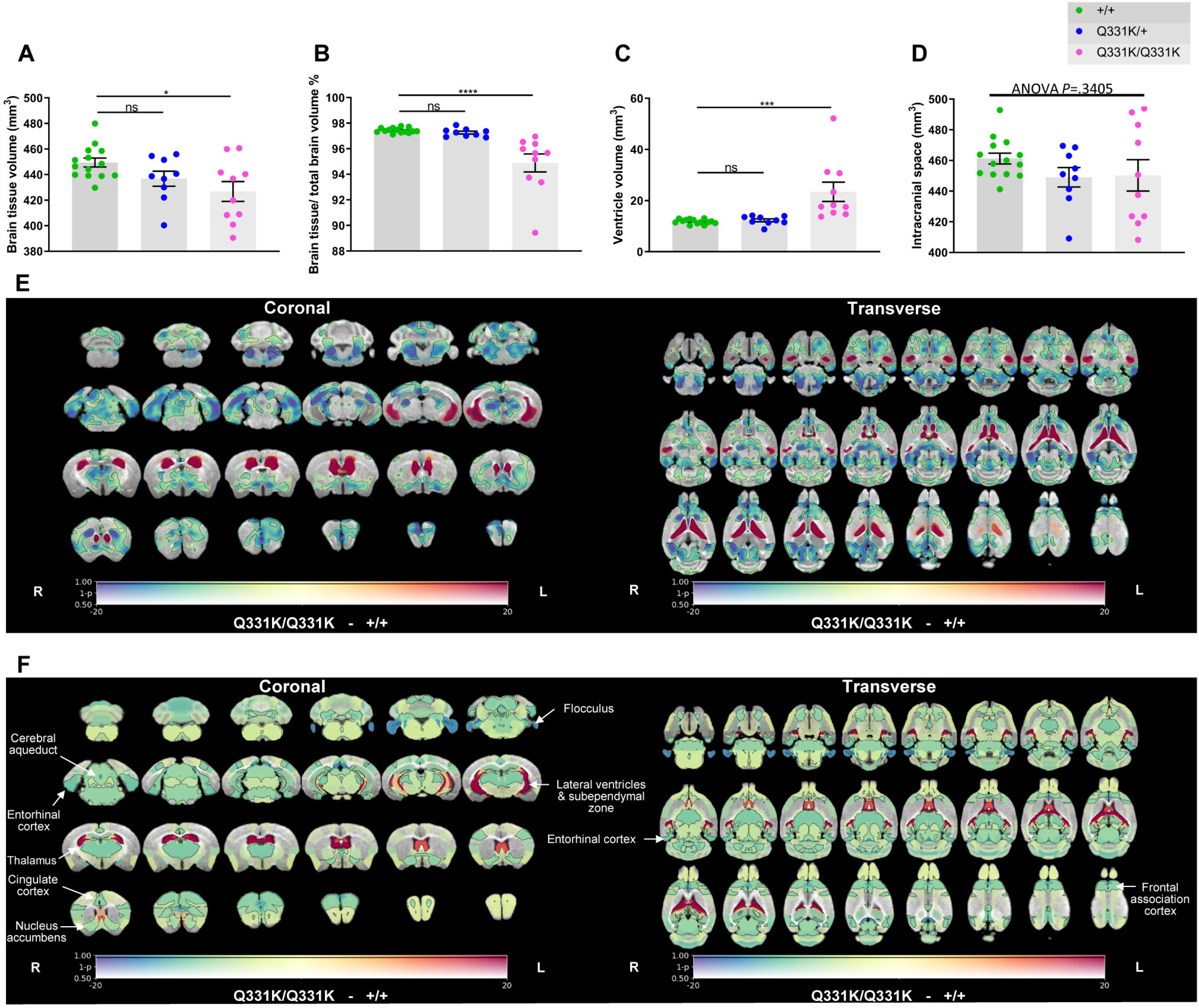
*in vivo* MRI reveals regional brain volume loss and ventricular enlargement in 7-month-old TDP-43^Q331K/Q331K^ knock-in mice. **A** Quantification of brain parenchyma volume. ANOVA *P*=0.0191. Pairwise comparisons: +/+ vs Q331K/+: ns *P*=0.1159; +/+ vs Q331K/Q331K: \**P*=0.0115. **B** Quantification of brain parenchyma volume as a percentage of total brain volume. ANOVA *P*<0.0001. Pairwise comparisons: +/+ vs Q331K/+: ns *P*=0.7448; +/+ vs Q331K/Q331K: \*\*\*\**P*<0.0001. **C** Quantification of ventricle volume. ANOVA *P*=0.0003. Pairwise comparisons: WT (+/+) and TDP-43^Q331K/+^ (Q331K/+): ns *P*=0.8597; WT (+/+) and TDP-43^Q331K/Q331K^ (Q331K/Q331K): \*\*\**P*=0.0004. **D** Quantification of brain parenchyma volume and ventricular volume combined. ANOVA *P*=0.3405. (**A-D**) Each dot represents one mouse. Groups were compared by one-way ANOVA followed by Holm-Sidak *post-hoc* test for pairwise comparisons; error bars represent mean ± s.e.m. **E** A map of voxelwise differences in volume between 7-month-old +/+ and Q331K/Q331K mice calculated from *in vivo* MR images, overlaid on the T1-weighted study-specific template. The map is displayed in the coronal plane (rostral-caudal) and the transverse plane (ventral-dorsal). The R and L indicate the right and left sides of the mouse. The colours of the overlay indicate the percent volume difference (cool colours indicate reduced volume in +/+ mice compared to Q331K/Q331K mice), and the opacity of the overlay indicates the significance of the volume difference (regions where the FWE-corrected *P*>0.5 are completely transparent, and regions where the FWE-corrected *P*=0 completely opaque). Black contours demarcate regions where the FWE-corrected *P*<0.05. **F** A similar map as in **E** but showing differences in the DSURQE atlas ROI volumes. Black contours demarcate ROIs where the FDR-corrected *P*<0.05. ROIs of particular interest are annotated.

To more precisely define patterns of atrophy and identify brain areas most vulnerable to mutant TDP-43^Q331K^, we conducted a region of interest (ROI) analysis using an anatomically parcellated mouse brain atlas (DSURQE, Mouse Imaging Centre, Toronto) (Dorr et al., 2008), which we registered to the MRI scans. This allowed us to compare the volumes of 182 predefined regions between the groups of mice. Fifty-one regions with statistically significant volume changes between TDP-43^Q331K/Q331K^ and wild-type mice were identified after correction for multiple comparisons at 5% false discovery rate (FDR) (Table 1 and Fig. 1F). The most significant cortical volume reductions occurred in the frontal, entorhinal, orbital and cingulate cortices, while subcortical volume loss was most marked in the nucleus accumbens, thalamus, subiculum, hippocampus, basal forebrain and the brain stem. In the cerebellum, there were decreases in the flocculus and the deep nuclei (dentate and interpositus), as well as in the cerebro-cerebellar white matter tracts (cerebellar peduncles). The most significant volume increase was, as expected, in the lateral ventricles, although the volumes of the third and fourth ventricles were not significantly affected and the cerebral aqueduct was decreased in size (Table 1). We also detected apparent volume increases in several grey and white matter ROIs, notably the CA3 hippocampal field and fimbria. However, these are likely to be artefacts caused by the proximity of these ROIs to the enlarged lateral ventricles, which can cause signal contamination due to partial volume effects, imperfect normalisation and smoothing used in TBM analysis (Manera et al., 2019). Overall, the pattern of regional brain volume reduction in TDP-43^Q331K/Q331K^ mice was reminiscent of that seen in patients with ALS-FTD and in presymptomatic carriers of ALS-FTD-linked gene mutations (Table 1).

**Table 1.**
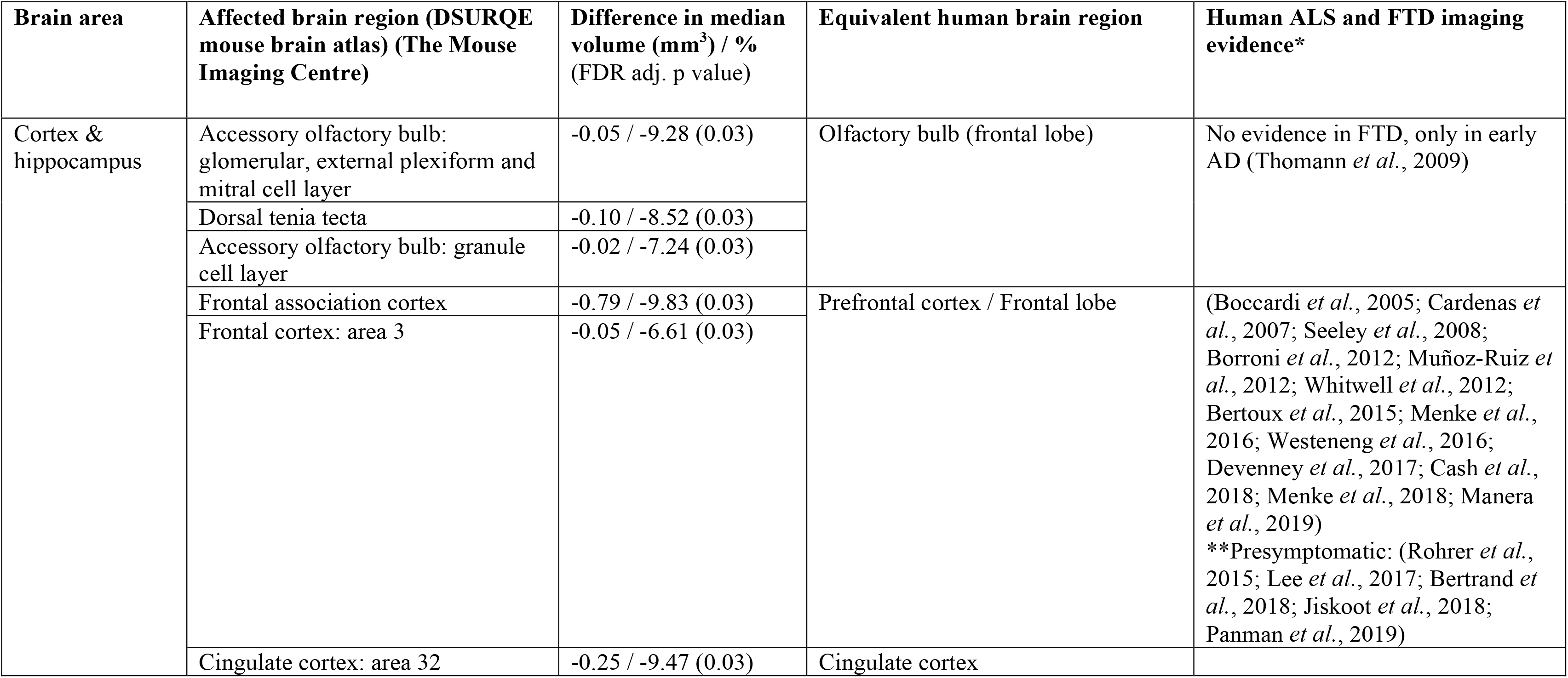
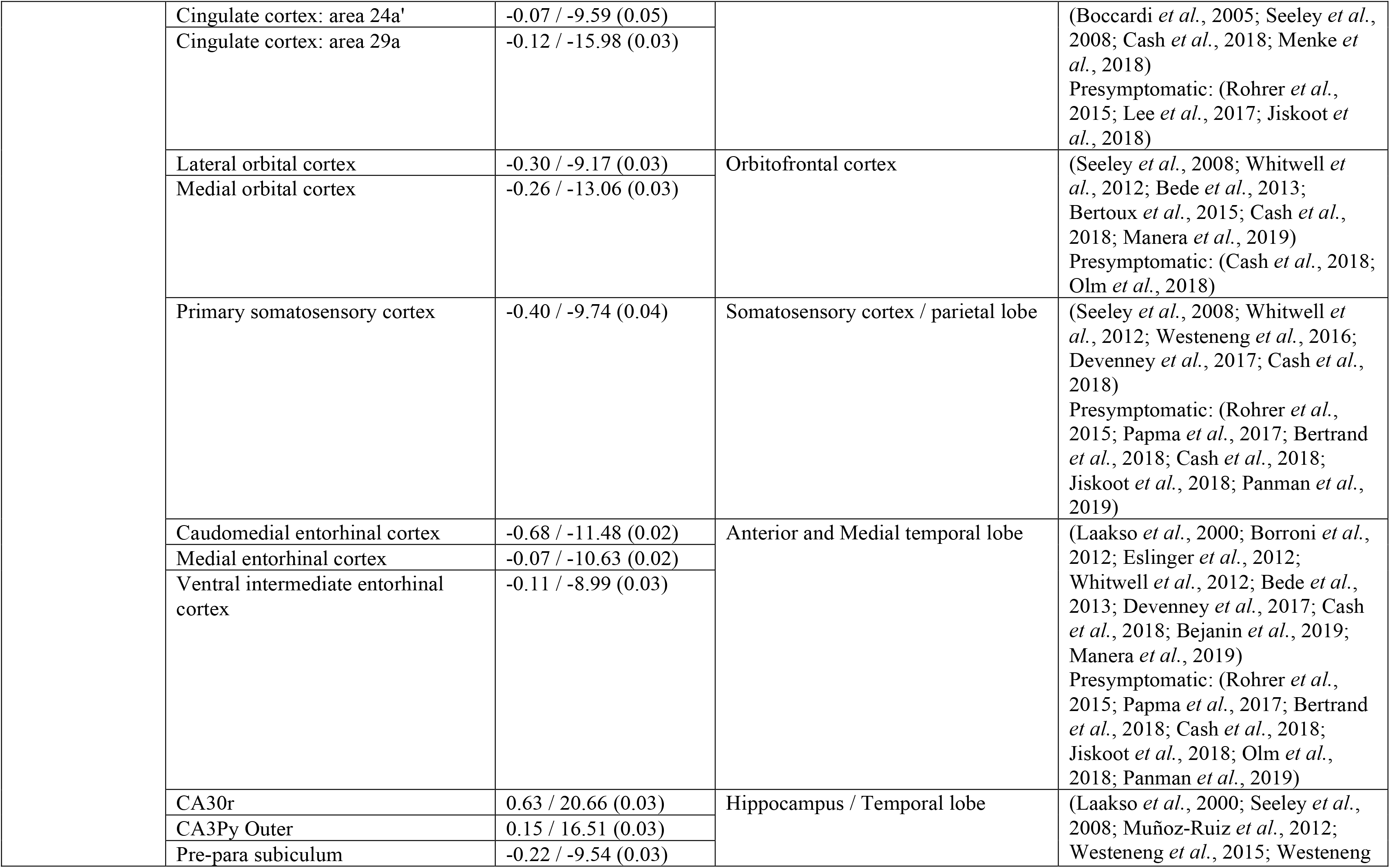
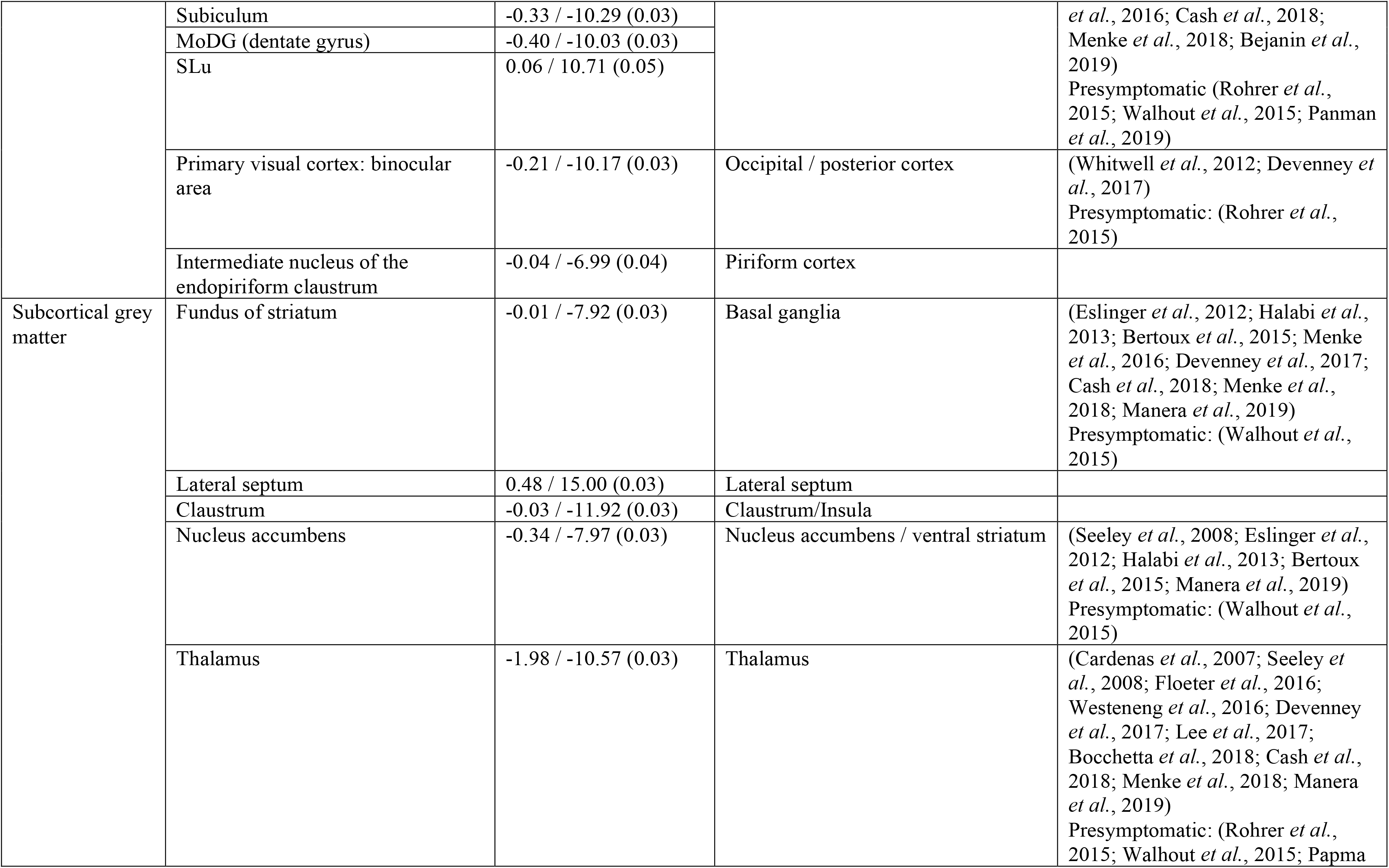
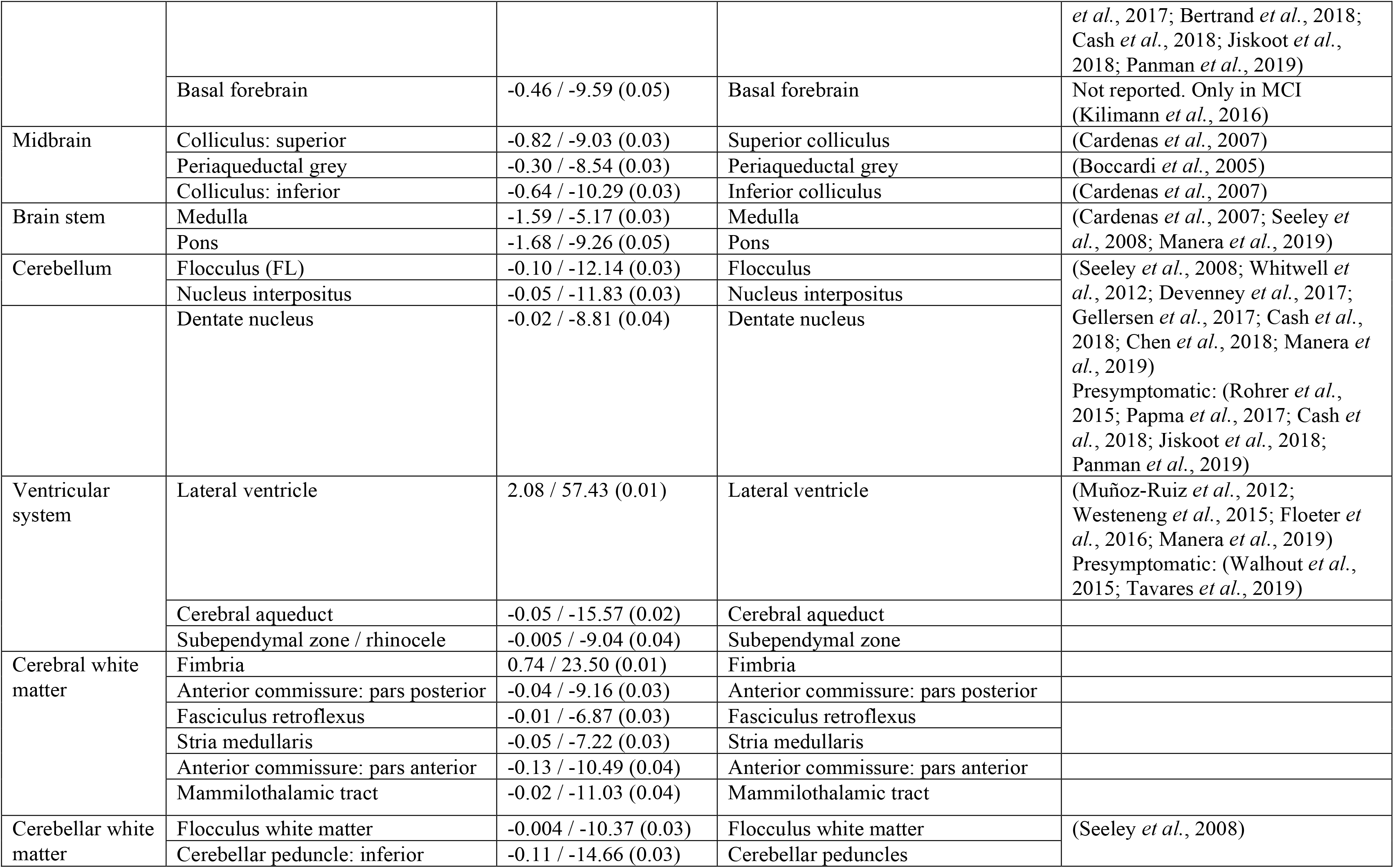
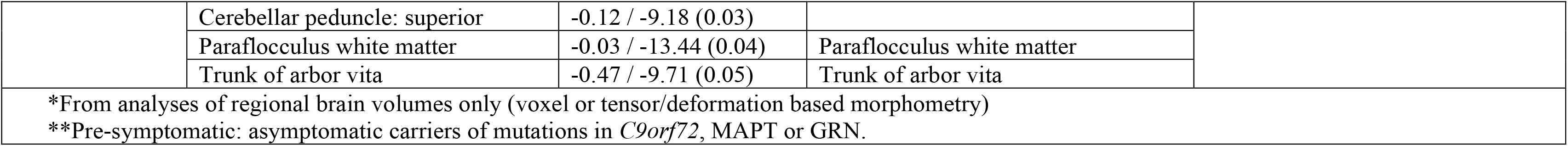
Structures with median volume (mm3) and % difference between TDP-43^Q331K/Q331K^ and wildtype mice that are significant at p<0.05 after FDR correction, and the corresponding or equivalent human brain areas as well evidence of similar/related changes reported in human ALS and FTD imaging literature

Comparison of heterozygous TDP-43^Q331K/+^ with wild-type mice yielded no statistically significant volume differences after correction for multiple comparisons at 5% FDR. However, TDP-43^Q331K/+^ mice did demonstrate trends towards volume loss or gain in several brain regions that were significant in TDP-43^Q331K/Q331K^ mice (Supplementary Figure 1A, B). This suggests a dose-effect of the mutation on brain structural phenotypes. Indeed, we made similar observations in our previous study with TDP-43^Q331K/+^ mice demonstrating similar phenotypes and transcriptomic changes to those seen in TDP-43^Q331K/Q331K^ mice, but of smaller magnitude (White et al., 2018). We therefore focused our attention on comparing wild-type with TDP-43^Q331K/Q331K^ mice in subsequent histological analyses in order to identify cellular correlates of volume loss on MRI.

### Reduced parvalbumin interneuron density in ROIs

We previously observed a reduced density of parvalbumin-positive (PV+) interneurons in frontal cortices of TDP-43^Q331K/Q331K^ mice as well as reduced PV mRNA expression at both 5 and 20 months of age (White et al., 2018). We reasoned that interneuron readouts could be a sensitive indicator of early disease in ALS-FTD. We therefore immunostained for PV in four ROIs demonstrating volume loss on MRI. We also analysed PV+ interneuron density in the visual cortex as a negative control as this area was largely unaffected on MRI (Fig. 2A, B). We found that the frontal cortex showed a reduction (by 27.1%) in PV+ interneuron density in mutant mice. The cingulate cortex was one of the most significantly affected subregions within the frontal cortex, demonstrating a 50.1% reduction of PV+ interneuron density. The entorhinal cortex and the dentate gyrus of the hippocampus demonstrated 60.2% and 39.8% reductions in PV+ interneuron density, respectively. By contrast, there was no significant reduction in PV+ interneuron density in the visual cortex (Fig. 2C). Finally, to determine if other interneuron subtypes could be affected by mutant TDP-43^Q331K^ we quantified somatostatin-positive (SOM+) interneurons. No significant change in SOM+ interneuron density was seen in mutant mice (Supplementary Fig. 2A, B). Collectively, these results indicate that PV+ interneurons are selectively reduced in number in MRI regions that are significantly reduced in volume, but not in an area that was not atrophic.

**Figure 2.**
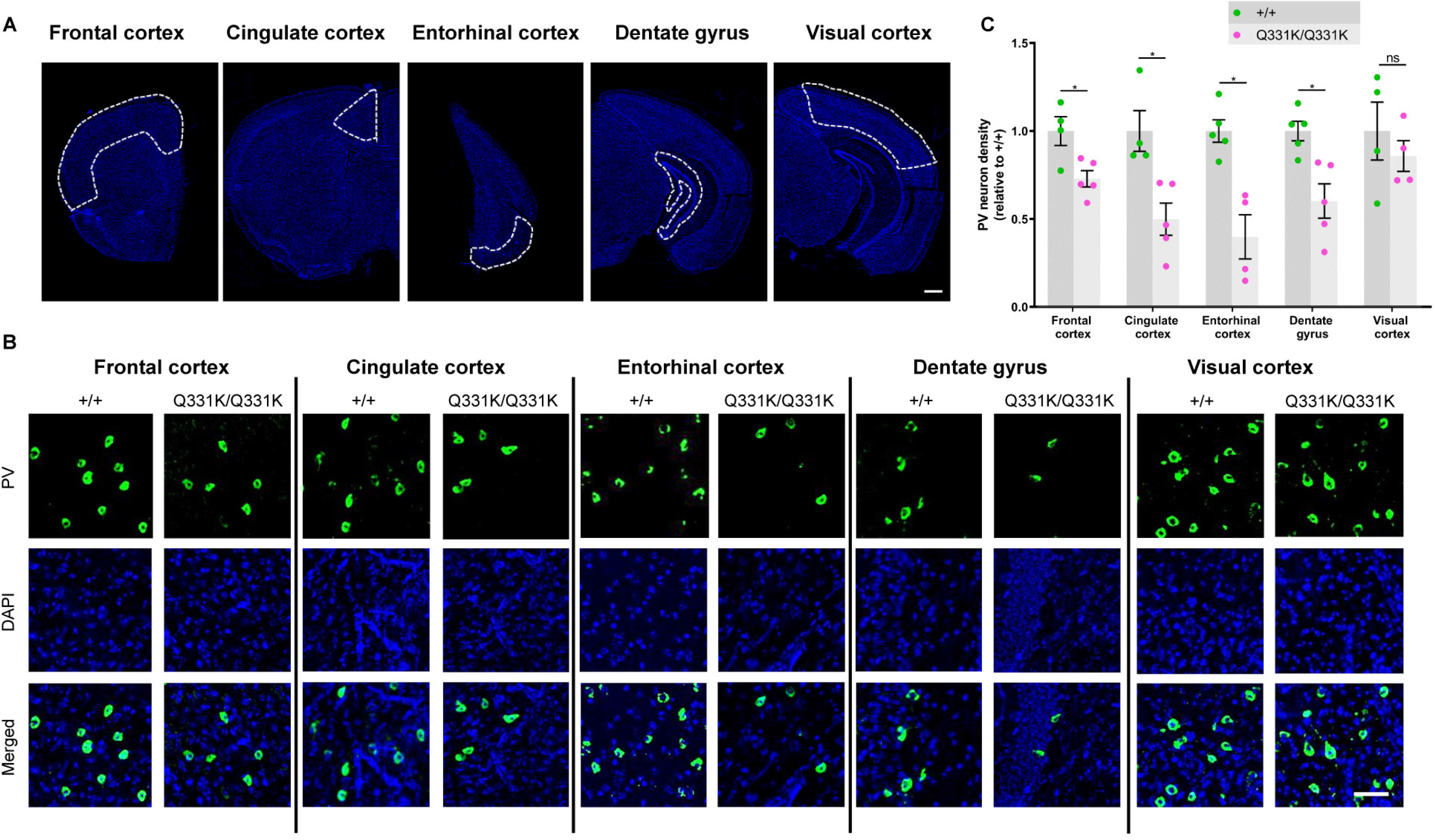
Reduced parvalbumin interneuron density in MRI regions of interest in 7-month-old TDP-43^Q331K/Q331K^ mice. **A** Low magnification DAPI-stained overview of indicated brain regions of 7-month-old mice. Scale bar, 500 μm. **B** Representative images showing parvalbumin (PV) staining and DAPI (blue) in given brain regions of +/+ and Q331K/Q331K mice. Scale bar, 50μm. **C** Quantification of PV+ interneuron density in given brain regions. Comparisons between +/+ and Q331K/Q331K as follows: frontal cortex, \**P*=0.0368; cingulate cortex, \**P*=0.0319; entorhinal cortex, \**P*=0.0129; dentate gyrus, \**P*=0.0294; visual cortex, ns *P*=0.4742. Each dot represents one mouse. *P* values were calculated with multiple t-tests adjusted by Holm-Sidak correction. Error bars represent mean ± s.e.m.

### Widespread microglial activation

Glial involvement in the pathogenesis of ALS-FTD is well established, with both astrocytic and microglial activation reported, while the presence of TDP-43 pathology in white matter implicates oligodendrocytes as well (Neumann *et al.*, 2007; Radford *et al.*, 2015; Umoh *et al.*, 2018). To determine the responses of glia to mutant TDP-43^Q331K^ we used immunohistochemistry to study two MRI-defined brain ROIs, and again used the visual cortex as a negative control region. We assessed microglia by staining for ionized calcium-binding adapter molecule 1 (Iba-1), which is elevated in activated microglia, and transmembrane protein 119 (Tmem-119), which is downregulated in disease-associated microglia (DAM) (Butovsky *et al.*, 2014; Deczkowska *et al.*, 2018). Iba1 immunostaining revealed significant increases in the area of staining and in microglial density in the frontal and entorhinal cortices of TDP-43^Q331K/Q331K^ mice, while Tmem-119 immunostaining was significantly reduced in these regions (Fig. 3A-D). Surprisingly, the visual cortex also displayed both increased Iba1 and decreased Tmem119 staining. Astrocytes, however, demonstrated no evidence of activation or proliferation in mutant mice by immunostaining for glial fibrillary acidic protein (GFAP) (Supplementary Fig. 3A, B). Immunostaining for myelin basic protein also showed no significant difference between mutant and wild-type mice, suggesting no change in myelination due to mutant TDP-43^Q331K^ (Supplementary Fig. 3C, D). Collectively, these results indicate that the earliest glial change in TDP-43^Q331K^ knock-in mice is the transformation of microglia into a disease-associated state, and that this occurs even in a region without volume loss on MRI.

**Figure 3.**
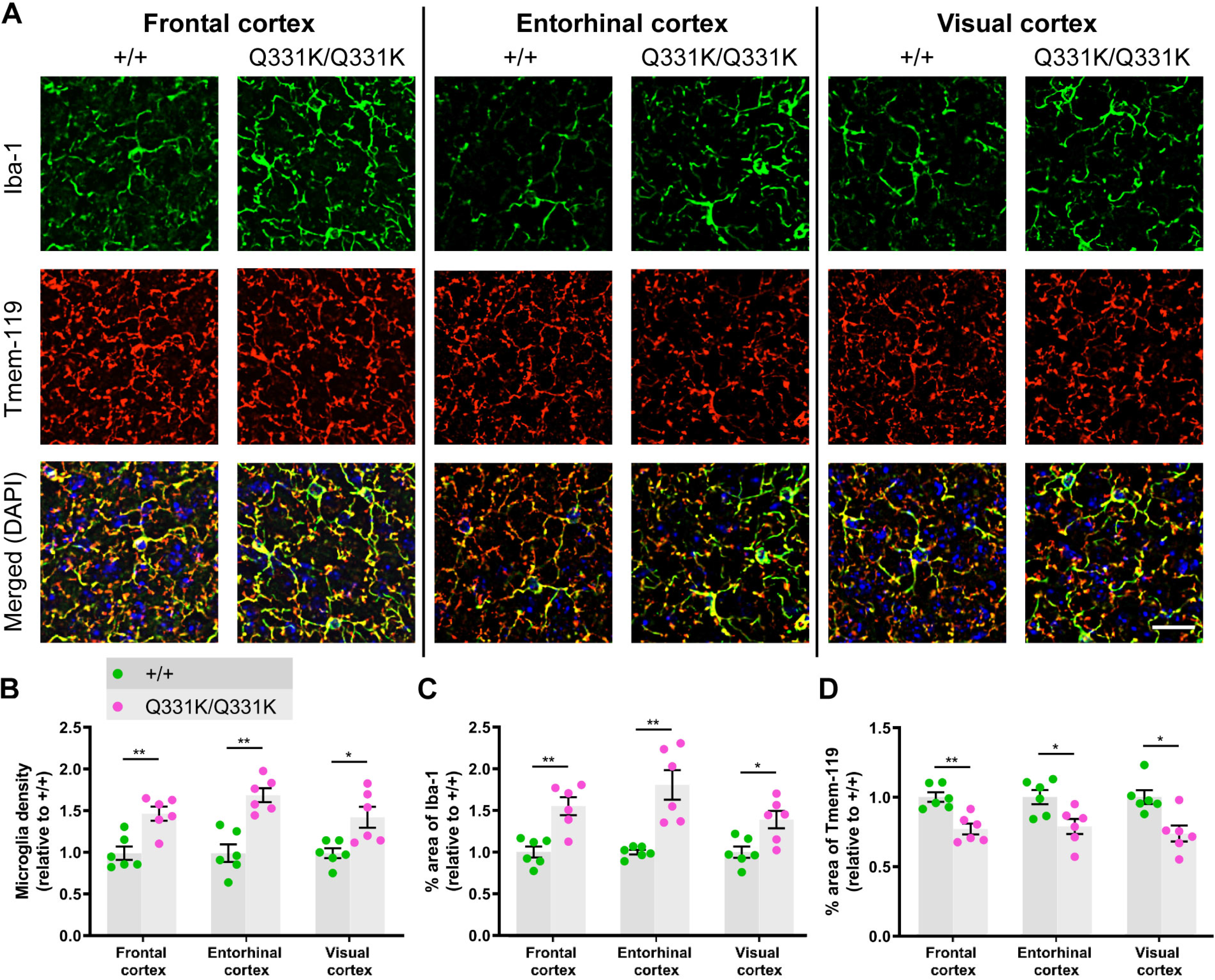
Global microglial activation in the TDP-43^Q331K/Q331K^ mutant mouse brain. **A** Immunohistochemistry showing microglia marker Iba-1 (green), Tmem-119 (red), and DAPI (blue) in indicated regions of 7-month-old +/+ and Q331K/Q331K mouse brains. Scale bar, 20μm. **B** Quantification of microglia density based on Iba-1 and DAPI immunoreactivity in given brain regions. Comparisons between +/+ and Q331K/Q331K as follows: frontal cortex, \*\**P*=0.0028; entorhinal cortex, \*\**P*=0.0012; visual cortex, \**P*=0.0113. **C** Quantification of percentage area of Iba-1 in given brain regions. Comparisons between +/+ and Q331K/Q331K: frontal cortex, \*\**P*=0.0043; entorhinal cortex, \*\**P*=0.0035; visual cortex, \**P*=0.0107. **D** Quantification of percentage area of Tmem-119 in given brain regions. Comparisons between +/+ and Q331K/Q331K: frontal cortex, \*\**P*=0.0039; entorhinal cortex, \**P*=0.0174; visual cortex, \**P*=0.0122. (**B-D**) Each dot represents one mouse. *P* values were calculated with multiple t-tests adjusted by Holm-Sidak correction. Error bars represent mean ± s.e.m.

### Reduced hippocampal neurogenesis

Another region demonstrating significant volume loss on MRI and reduced PV+ interneuron density was the hippocampal dentate gyrus (DG). Given that this structure is an important niche for adult hippocampal neurogenesis (AHN) and that PV+ interneurons have an essential role in neurogenesis (Song et al., 2013; Groisman et al., 2020) we quantified markers of cell proliferation (Ki67) and immature neuron formation (doublecortin, DCX) in the granule cell layer (GCL) of the DG. We found no difference in the number of Ki67+ cells between wildtype and TDP-43^Q331K/Q331K^ mice, suggesting that the rate of progenitor cell division was unaffected (Fig. 4A, B). However, quantification of DCX+ cells revealed a significant reduction in the number of immature neurons in the GCL of TDP-43^Q331K/Q331K^ mice (Fig. 4A, C). Given the volume changes in the lateral ventricles we also examined the subventricular zone (SVZ), another key area of adult neurogenesis, but found no evidence of impaired neurogenesis here (Fig. 4D-F). Collectively, these results indicate that mutant TDP-43^Q331K^ causes a reduction in the number of progenitor cells undergoing neuronal differentiation or, alternatively, reduces the number of cells surviving to the immature neuron stage in the DG, and that a paucity of PV+ interneurons could be contributing to this.

### Aberrant neurodevelopment

One of the most striking observations on MRI was the lateral ventricular enlargement in TDP-43^Q331K/Q331K^ mice. This could simply be secondary to a reduction in the brain parenchyma, but another explanation is that the ventricular system had not developed properly and that the ventriculomegaly actually represents a state of hydrocephalus. Indeed, the cerebral aqueduct was significantly reduced in volume in mutants (Table 1), indicating that CSF outflow could be compromised. Furthermore, occasional juvenile mutants demonstrated frank hydrocephalus with doming of the skull and had to be euthanised before 6 weeks of age (Figure 5A, B). We postulated that the ventricular enlargement of mutants that survived through to later adulthood could also have occurred early in life, before significant ossification of cranial sutures, and that this would be reflected as a change in skull shape that would persist into adulthood once skull plates had fused. We therefore performed *ex vivo* micro computerised tomography (CT) on heads of aged 24-month-old male mice. Indeed, geometric morphometric comparison of TDP-43^Q331K/Q331K^ and wild-type littermate skulls using principal component analysis revealed a significant difference in shape between the two genotypes (Figure 5C, D). Specifically, a subtly dysmorphic craniofacial structure was seen in mutants characterised by midfacial hypoplasia and brachycephaly – effectively an increase in skull sphericity – and a reduction in overall skull size (Figure 5E, Supplementary video 1). These changes are in keeping with the hypothesis that TDP-43^Q331K^ disrupts the development of the ventricular system.

**Figure 4.**
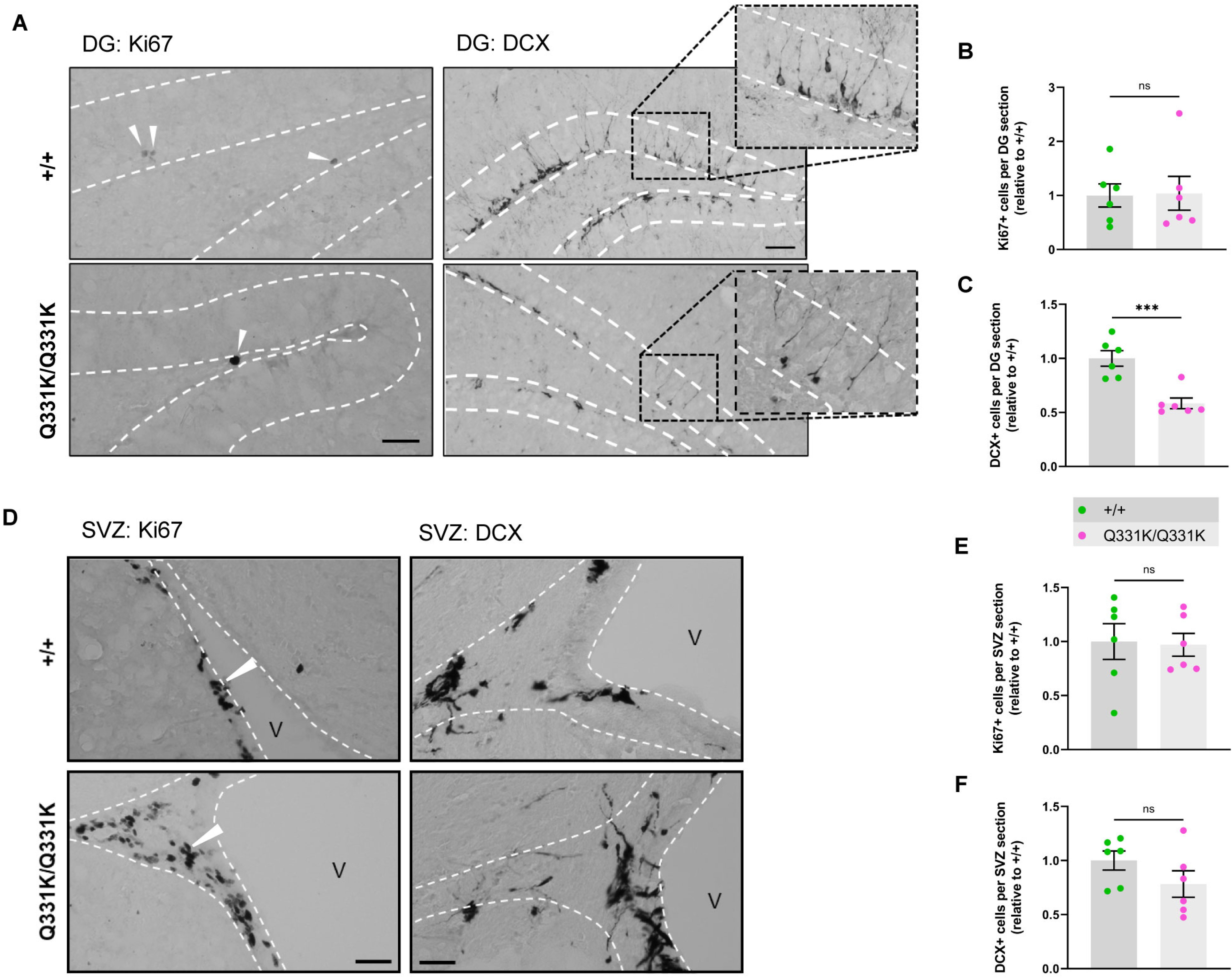
Aberrant neurogenesis in TDP-43^Q331K/Q331K^ mice. **A** Immunohistochemistry showing cell division marker Ki67 (arrow heads) and immature neuron marker DCX in the mouse hippocampal dentate gyrus of 7-month-old +/+ and Q331K/Q331K mice. Scale bar, 40 and 100 μm, representatively. **B** Quantification of Ki67+ cells per dental gyrus (DG) section in hippocampus between +/+ and Q331K/Q331K mice: ns *P*=0.9183. **C** Quantification of DCX+ cells per DG section in the hippocampal dentate gyrus between +/+ and Q331K/Q331K mice: \*\*\**P*=0.0008. **D** Representative images showing cell division marker Ki67 (arrow heads) and immature neuron marker DCX in the mouse subventricular zone with **E, F** quantification. Comparisons between +/+ and Q331K/Q331K as follows: Ki67, ns *P*=0.8820; DCX, ns *P*=0.1794. V = lateral ventricle. (**B, C, E, F**) Each dot represents one mouse. Groups were compared by unpaired 2-tailed t-test All data shown are mean ± s.e.m.

**Figure 5.**
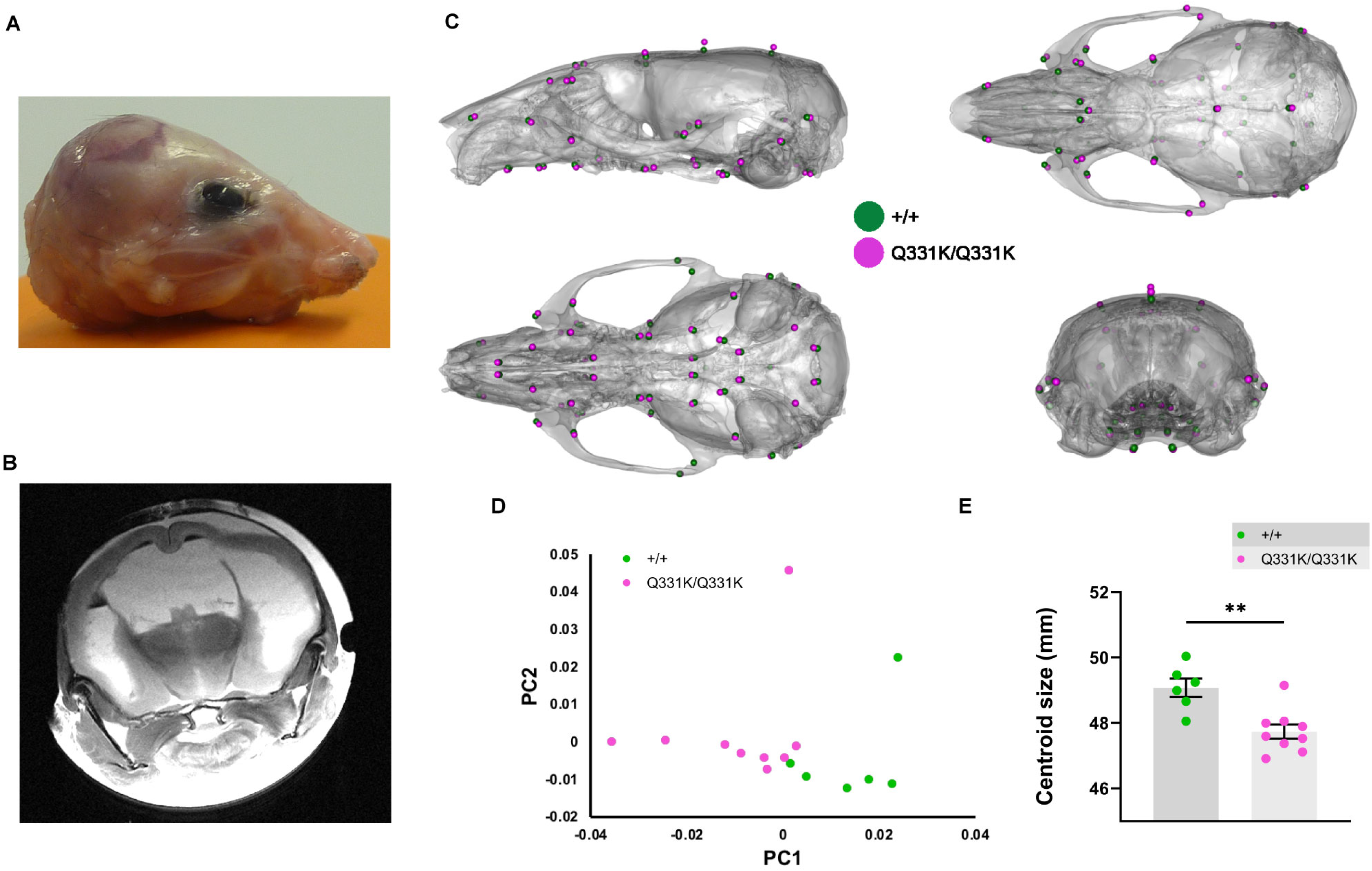
Altered skull shape in TDP-43^Q331K/Q331K^ mice. **A** Photo of scalped head of a Q331K/Q331K mouse and **B** *ex vivo* MRI scan of fixed head of the mouse. **C** Geometric morphometric analysis of +/+ vs Q331K/Q331K mice. Average landmark configurations of +/+ and Q331K/Q331K skulls superimposed on +/+ average skull, showing (clockwise from top left) lateral, superior, rear and inferior views. Each dot represents a single standardised landmark, the colour indicating the average for each genotype. **D** Principal component analysis (first two components) of skull shape variation after scaling. **E** Centroid sizes (the square root of the sum of the squared distances of a set of landmarks from their collective centroid) of +/+ and Q331K/Q331K skulls. Groups were compared by unpaired 2-tailed t-test: \*\**P*=0.0022. Error bars represent mean ± s.e.m. (**D, E**) Each dot represents one mouse.

We also wished to determine if mutant TDP-43^Q331K^ influenced the development of the brain parenchyma itself. We therefore turned to our histological readout of choice, PV+ interneuron density, looking specifically at the hippocampi of juvenile P14 mouse pups (Figure 6A). We found a significant reduction of PV+ interneuron density in the DG of mutant mice, particularly the hilus, but no changes in the CA3 region (Figure 6B-E). These observations establish a role for TDP-43 in the development of PV+ interneurons in the brain, which in turn suggests that the paucity of PV+ interneurons in adult mice is at least partly due to their failure to develop in the first place.

**Figure 6.**
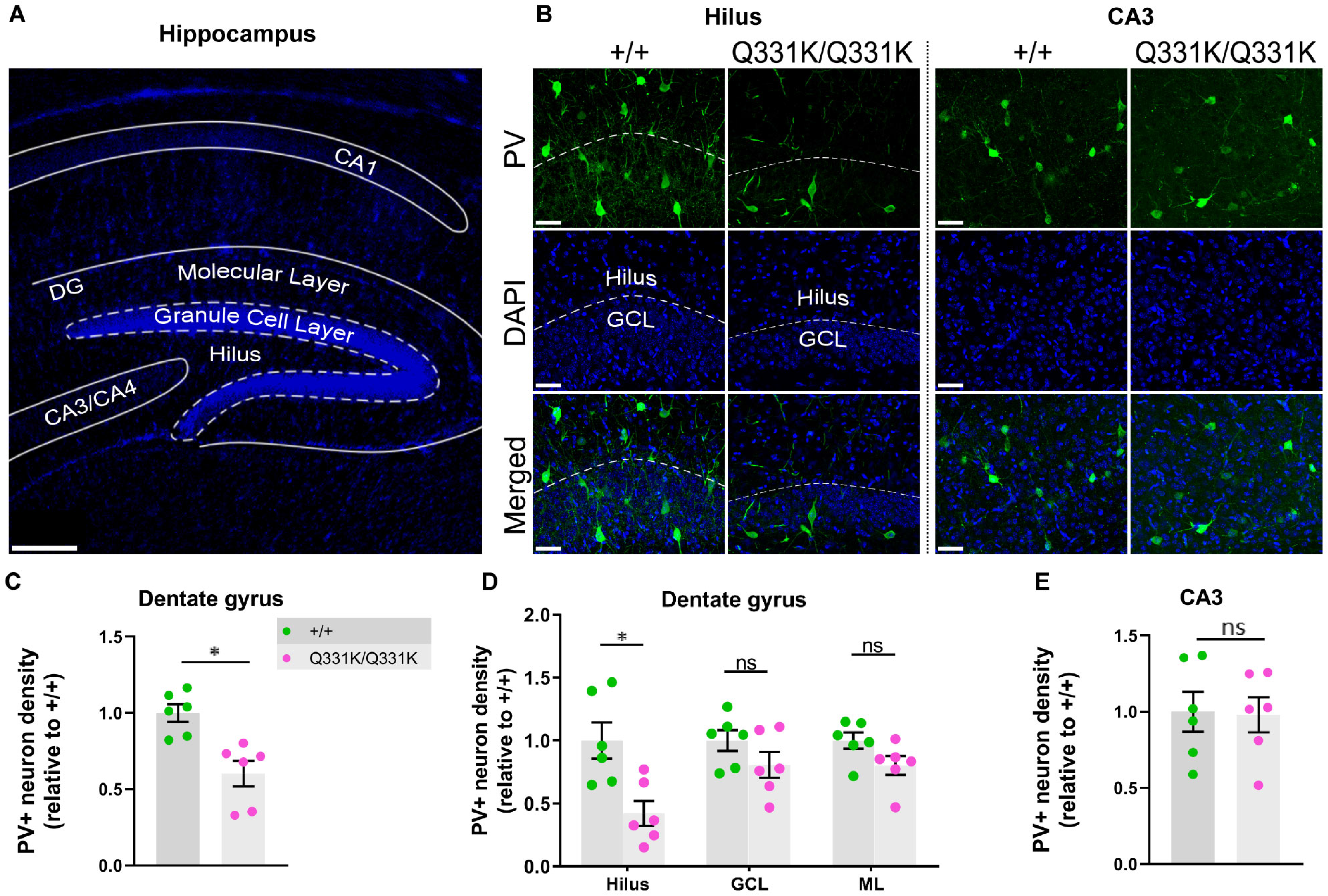
Paucity of PV+ neurons in hippocampi of P14 TDP-43^Q331K/Q331K^ mice. **A** Low magnification DAPI-stained overview of mouse P14 hippocampal regions. Scale bar, 200 μm. **B** PV+ interneurons shown in given regions of the hippocampus in P14 +/+ and Q331K/Q331K mice. Scale bar, 50 μm. **C** Quantification of PV+ interneuron density in dentate gyrus (DG) comparing +/+ to Q331K/Q331K mice: \**P*=0.0142. **D** Of regions within DG, quantification of PV+ interneuron density in hilus comparing +/+ to Q331K/Q331K mice: \**P*=0.0315; granule cell layer (GCL): ns *P*=0.3133 and molecular layer (ML): ns *P*=0.1957. **E** Quantification of PV+ interneuron density in C3 area: ns *P*=0.9061. (**C, D**) Each dot represents one mouse. *P* values were calculated with multiple t-tests adjusted by Holm-Sidak correction. Error bars represent mean ± s.e.m.

### Paucity of PV+ interneurons in human ALS

Our MRI-guided histological survey in mutant mice indicated a central role for PV+ interneurons in several aspects of TDP-43^Q331K^-mediated disease: regional vulnerability of the brain, aberrant neurogenesis and neurodevelopment. We therefore sought to determine if disruption to PV+ interneurons was a feature of human disease. We immunostained for PV in the dorsolateral prefrontal cortex of 18 patients with ALS patients (6 with *C9orf72* mutations and 12 sporadic ALS cases without known ALS gene mutations) and 11 age and sex-matched neurologically normal controls (Figure 7A). Demographic details of brain donors are given in Supplementary tables 2 and 3, including post mortem interval (PMI), sex, age and *C9orf72* genetic status. There was no significant effect of PMI or age on PV+ interneuron density (Figure 7B, C). However, we detected a 24.6% reduction of PV+ interneuron density in ALS patients relative to controls (Figure 7A, D) with both mutant *C9orf72*-linked ALS and sporadic ALS cases showing a similar deficiency. This confirms that a paucity of PV+ interneurons in the frontal cortex is a feature of human ALS.

**Figure 7.**
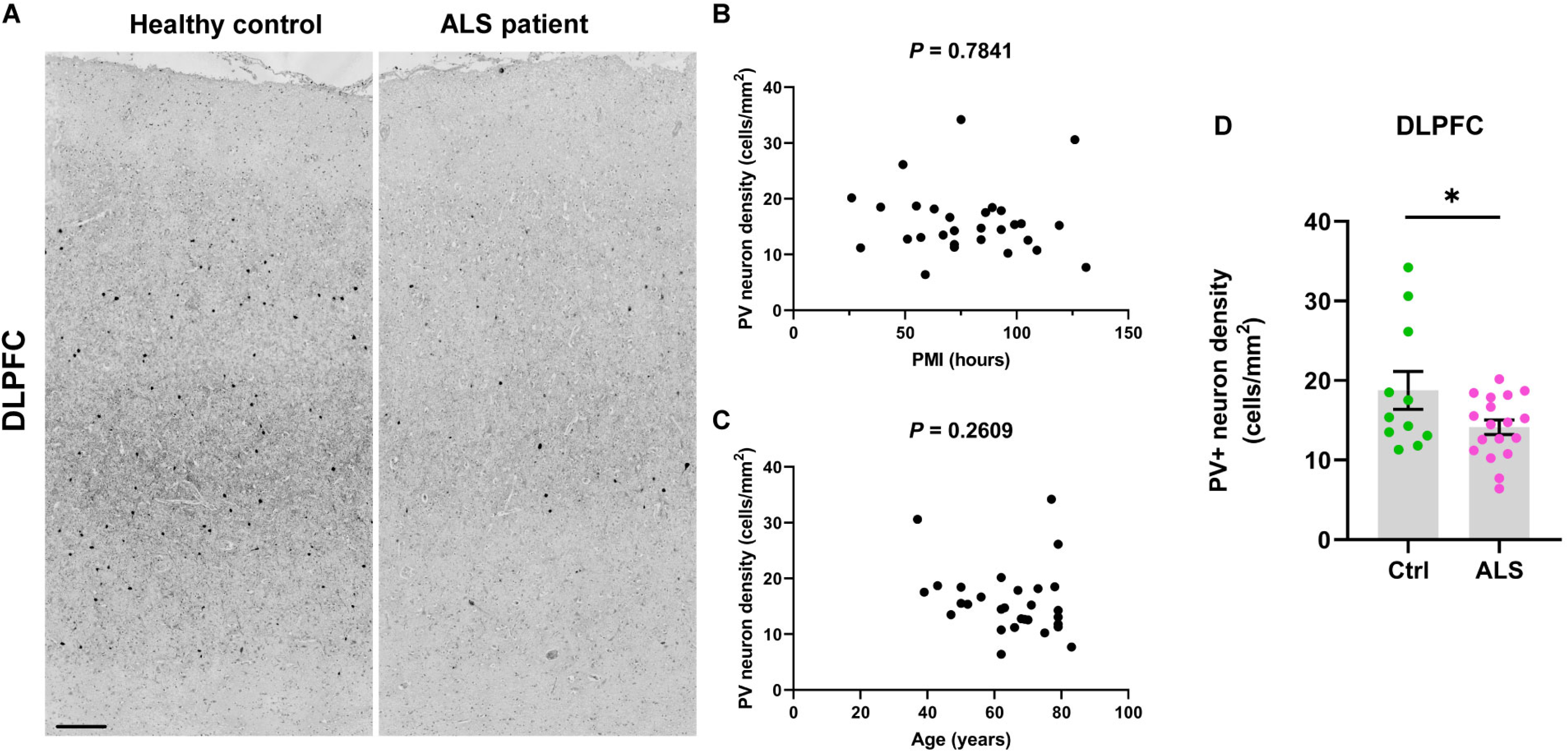
Paucity of PV+ neurons in DLPFC of ALS patients. **A** Immunohistochemistry showing PV+ interneurons in the dorsolateral prefrontal cortex (DLPFC) of ALS patients and neurologically healthy controls. Scale bar, 300 μm. **B** Scatter plot demonstrating PV+ interneuron density versus post mortem interval for each case (cases and controls combined). **C** Scatter plot demonstrating PV+ interneuron density versus age of each patient (cases and controls combined). **D** Quantification of PV+ interneuron density (cells/mm^2^) in DLPFC: * *P*=0.0447. Each dot represents one mouse. Groups were compared by unpaired 2-tailed t-test: Error bars represent mean ± s.e.m.

## Discussion

Treatment for neurodegenerative diseases is likely to be most effective during presymptomatic or prodromal stages, and biomarkers sensitive to these periods are needed. Preclinical models that accurately reflect human disease are therefore essential, facilitating the discovery of molecular pathological changes underlying the very earliest stages of disease. Here, using structural MRI, we detected an extensive pattern of brain volume loss in TDP-43^Q331K^ knockin mice, a preclinical model for ALS-FTD, while still at an early stage of disease. Significantly, many of the brain areas affected in mutant mice are analogous to those involved in patients with FTD (Meeter *et al.*, 2017; Bruun *et al.*, 2019). Similar regions have also been implicated in recent studies of patients with ALS (Menke *et al.*, 2017). Perhaps most importantly, pre-symptomatic carriers of ALS-FTD-linked gene mutations also demonstrate atrophy in these brain areas, even decades in advance of clinical onset (Rohrer *et al.*, 2015; Cash *et al.*, 2018; Panman *et al.*, 2019). A detailed list of affected mouse and human brain regions is given in Table 1. Collectively, these observations underscore the value of the TDP-43^Q331K^ knock-in mouse in revealing *in vivo* structural biomarkers relevant to the pathogenesis of disease in patients with ALS-FTD.

Interestingly, our MRI studies of TDP-43^Q331K^ mice highlighted numerous regions beyond the frontal and temporal lobes that have only relatively recently also emerged as being affected in patients. Thalamic regions were smaller in mutant mice, mirroring observations in patients with sporadic and familial FTD caused by mutations in *C9orf72*, *MAPT* and *GRN*. The thalamus represents an important hub, modulating the flow of information between the external environment and the cortex, and could be contributing to many facets of FTD including abnormalities of cognition, attention and personality. The cerebellum was also affected in mutant mice, and it has been shown that cerebellar atrophy occurs in both mutant *C9orf72* and *MAPT* associated FTD (Cash *et al.*, 2018). Furthermore, cerebellar integrity has been associated with cognitive performance (Chen *et al.*, 2018), while functional MRI has identified impaired connectivity of cerebellar regions with the cerebrum in FTD (Guo *et al.*, 2016). Dysfunctional cerebellar and cortico-cerebellar circuits, particularly involving the vermis and the deep nuclei, are recognized as core contributors to psychosis, which can be part of the clinical picture of FTD (Schmahmann, 2016). Finally, the flocculus, which functions in ocular motility, was also affected in mutant mice, and this is notable as MRI changes in the cerebellum and eye movement abnormalities can occur in patients with ALS (Prell and Grosskreutz, 2013; Kang *et al.*, 2018).

Enlargement of the lateral ventricles is also emerging as a sensitive MRI biomarker in human FTD (Manera *et al.*, 2019; Tavares *et al.*, 2019). This may be due to subcortical parenchymal volume loss since one of the imaging hallmarks of FTD, and a point of divergence from AD, appears to be greater subcortical than cortical brain atrophy (Landin-Romero *et al.*, 2017). It was therefore remarkable that TDP-43^Q331K^ mice also had enlarged lateral ventricles. While this ventriculomegaly could be due to loss of brain parenchyma, it could also reflect hydrocephalus. In this respect, it is notable that elderly patients with FTD and concomitant hydrocephalus can show clinical improvement following cerebrospinal fluid (CSF) removal (Korhonen *et al.*, 2017; Korhonen *et al.*, 2019). Assaying CSF pressure in TDP-43^Q331K^ mice would therefore be of interest. Nonetheless, our observation that frank hydrocephalus occurred in a small number of juvenile TDP-43^Q331K^ mice is enough to suggest that a developmental contribution towards ventricular enlargement in mutants is likely. Corroborating this, our micro-CT analysis elucidated that even aged TDP-43^Q331K^ mice have more spherical skulls, suggesting that intracranial pressure was raised prior to skull plate fusion (i.e. during development). Although skull shape changes can occur in humans in association with age-related brain atrophy (Urban *et al.*, 2016), we conclude that enlargement of ventricles in TDP-43^Q331K^ mice is at least partly developmental in origin.

The recapitulation in TDP-43^Q331K^ knock-in mice of analogous brain MRI changes seen in prodromal human ALS-FTD allowed us to address a fundamental question in neurology: how do disturbances in ubiquitously expressed proteins such as TDP-43 cause regionally selective degeneration in the nervous system? We approached this question by performing a histological survey and found that TDP-43^Q331K^ knock-in mice were deficient in PV+ interneurons in areas showing atrophy on MRI, but not in an area with no volume loss. This suggests that PV+ interneurons contribute to regional brain vulnerability in ALS-FTD. Indeed, we validated that PV+ interneurons are reduced in human ALS frontal cortex. Previous studies of cortical GABAergic interneurons in ALS have identified robust reductions in calbindin D-28 immunoreactive cells in the motor cortex (Ferrer *et al.*, 1993; Maekawa *et al.*, 2004), but analyses of PV+ interneuron numbers have been less conclusive. One study found a significant decrease of PV+ interneurons in the motor cortex but did not examine more anterior regions of the brain (Nihei *et al.*, 1993), while another found only a trend towards a reduction in layer VI of the frontal cortex (Maekawa 2004). A third study found no decrease in PV+ interneurons in the frontal cortex in two patients with ALS with dementia (Ferrer *et al.*, 1993). Our study more strongly points towards a reduction in PV+ interneurons as being correlated with ALS and, furthermore, shows that both sALS and *C9orf72*-linked ALS display a similar degree of loss. In theory, such a loss of PV+ interneurons would lead to a reduction in cortical inhibition, which could have excitotoxic consequences for the pyramidal projection neurons onto which they synapse (Kiernan *et al.*, 2019). In support of this proposal, patients with mutations in *C9orf72*, the most common genetic cause of ALS and FTD, have been found to demonstrate cortical hyperexcitability (Geevasinga *et al.*, 2014). Thus, promoting PV+ interneuron activity is worthy of further study as a possible disease-modifying approach.

Intriguingly, we also observed a paucity of PV+ interneurons in P14 mice. This suggests that the deficiency of these cells in adult mice may be because they did not develop in the first place. Fast-spiking PV+ interneurons are the main modulators of synaptic plasticity during the critical period of cortical development, and disruption of interneurons leads to abnormalities in neuronal excitability, a feature of neurodevelopmental disorders, such as autism and schizophrenia (Hu *et al.*, 2014; Marin, 2016; Tremblay *et al.*, 2016). While ALS-FTD is not regarded as a developmental disease, mutations in some ALS-causing genes have been associated with juvenile-onset ALS with learning difficulties (Picher-Martel *et al.*, 2020), and studies have suggested that network degeneration in asymptomatic carriers of *C9orf72* mutations could be a consequence of aberrant patterning during development (Lee *et al.*, 2017). Thus, it is not inconceivable that disturbances in cortical excitation-inhibition balance during development may later predispose an individual to developing an age-related neurodegenerative disease.

Another important role served by PV+ interneurons is that of promoting hippocampal neurogenesis in the adult brain (Song *et al.*, 2013). We were therefore intrigued by the observation that the DG of the hippocampus, a niche for AHN, was smaller in TDP-43^Q331K^ mice and demonstrated a significant loss of PV+ interneurons. Examination of the number of immature neurons confirmed that AHN was indeed impaired in mutant mice. While AHN has been shown to be impaired in Alzheimer’s disease (AD) (Choi *et al.*, 2018; Moreno-Jimenez *et al.*, 2019), neurogenesis has been little studied in ALS-FTD. Interestingly, however, one human post-mortem study identified a reduction in the proliferation of neural progenitor cells in the hippocampi of patients with ALS (Galan *et al.*, 2017). Furthermore, overexpression of TDP-43 in neural progenitor cells of the developing mouse telencephalon was found to impair neurogenesis (Vogt *et al.*, 2018), while knockdown of TDP-43 *in vitro* resulted in increased expression of neurogenin 2, a transcription factor that specifies neuronal fate during development (Di Carlo *et al.*, 2013). Our study adds to this literature by demonstrating an *in vivo* role for TDP-43 in neurogenesis within the adult brain. Given the importance of adult-born hippocampal neurons to spatial memory, pattern separation, mood and cognition (Sahay *et al.*, 2011; Luna *et al.*, 2019), impaired AHN due to TDP-43 misregulation may well have contributed to the behavioural phenotypes we previously described in mutant mice (White *et al.*, 2018). Further studies to delineate the link between PV+ interneurons and AHN are warranted as we speculate that this will help find ways to leverage the neurogenic potential of the brain for therapeutic benefit in ALS-FTD.

In our analysis of non-neuronal cells we noted that microglia were in an activated state in TDP-43^Q331K^ knock-in mice. Microglia are thought to have a ‘surveying’ role when quiescent and become activated in neurodegenerative diseases, engulfing cellular debris, phagocytosing protein aggregates and pruning synapses, possibly through complement activation (Wake *et al.*, 2013; Hong *et al.*, 2016; Paolicelli *et al.*, 2017). Using inducible TDP-43 transgenic mice it has been shown that microglia play a key role in recovery from neurodegeneration (Spiller *et al.*, 2018). Unexpectedly, we found that microglial activation occurred even in the visual cortex, an area that was largely unaffected on MRI and which was not deficient in PV+ interneurons. This suggests that microglial activation is widespread and does not explain regional brain vulnerability in TDP-43^Q331K^ knock-in mice, at least when 7 months old. Although activated microglia in the mouse brain are regionally and functionally heterogeneous, and they may not be acting destructively, the reduced Tmem119 we observed is indicative of a shift to a DAM phenotype in mutants. Further work is needed to examine the causes and consequences of this microglial activation.

In conclusion, MRI changes in human ALS-FTD are recapitulated in TDP-43^Q331K^ knock-in mice, allowing this model to be used to understand the cellular and molecular meaning of macroscopic brain changes, something that is currently impossible in humans. Further studies of TDP-43^Q331K^ knock-in mice using longitudinal structural and functional MRI and MR spectroscopy promises to help delineate the natural history of ALS-FTD. More detailed examination of molecular changes within affected and unaffected brain regions, for example using *in situ* transcriptomic approaches, should help to unravel how different neuronal and glial subtypes contribute to disease, including during development and in neurogenic niches. These profiles will ultimately be of value in measuring the efficacy of therapeutic approaches designed to treat and perhaps even prevent ALS-FTD.

## Acknowledgements

We thank Babraham Institute Experimental Unit staff for technical assistance, Dr. George Chennell of the Wohl Cellular Imaging Centre at Maurice Wohl Clinical Neuroscience Institute, King’s College London, for assisting with image acquisition and analysis. We thank Dr Marta Vicente Rodrigues of the BRAIN Centre, Department of Neuroimaging, IoPPN, King’s College London, for assisting with immunohistochemistry. We acknowledge the provision of human brain tissues from the MRC Edinburgh Brain & Tissue Bank.

## Supplementary Material

**Supplementary fig. 1** No significant volume change in ROI analysis comparing heterozygous mutant to wild-type mice.

**Supplementary fig. 2** No significant change in SOM+ interneuron density in frontal and cingulate cortices in mutant mice.

**Supplementary fig. 3** No significant changes in astrocyte density and MBP % area in TDP-43^Q331K^ mice.

**Supplementary video 1** Skull morph between representative wild-type and TDP-43^Q331K/Q331K^ skulls, where size has not been regressed out.

**Supplementary table 1** Antibodies used in immunohistochemistry.

**Supplementary table 2** Patients’ MRC numbers and brain bank conditions.

**Supplementary table 3** Controls’ MRC numbers and brain bank conditions.

