## Supplementary Materials for "MRI-guided histology of TDP-43 knock-in mice implicates parvalbumin interneuron loss, impaired neurogenesis and aberrant neurodevelopment in ALS-FTD"

A

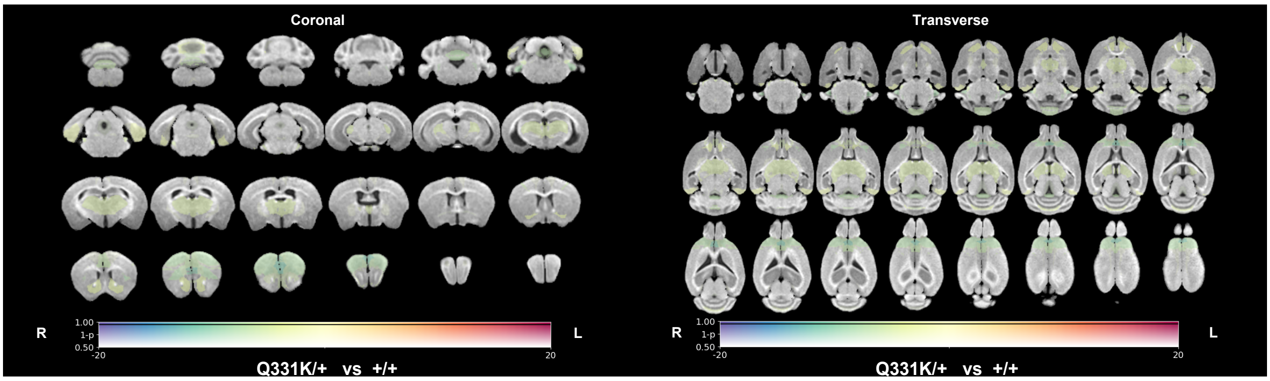


B

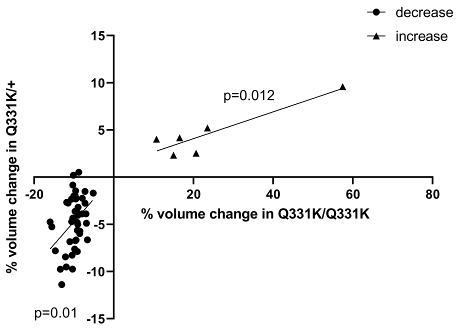


**Supplementary fig. 1 No significant volume change in ROI analysis comparing heterozygous mutant to wild-type mice. A** MRI study-specific template (coronal and transverse) with an overlay representing ROI volume differences at 7 months of age between +/+ and Q331K/+ mice. The colour of the overlay indicates the inter-group volume difference, while the transparency indicates the statistical significance. ROIs in which FWE-corrected *P*<0.05 are contoured in black. **B** Correlation between % volume changes in ROIs in Q331K/+ and Q331K/Q331K mice compared to +/+ mice

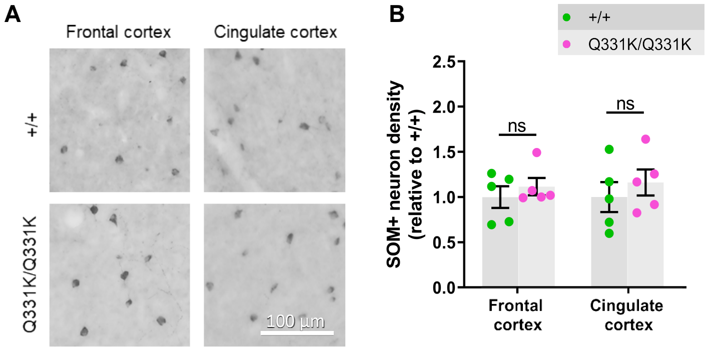


**Supplementary fig. 2 No significant change in SOM+ interneuron density in frontal and cingulate cortices in mutant mice. A** Representative images showing somatostatin positive (SOM+) neuron staining (dark brown) in frontal and cingulate cortices of 7-month-old +/+ and Q331K/Q331K mice. **B** Quantification of SOM+ neurons density in frontal cortex, ns *P*= 0.7204; cingulate cortex, ns *P*=0.7204 in 7-month-old Q331K/Q331K mice compared to +/+. *P* values were calculated with multiple t-tests adjusted by Holm-Sidak correction. n=5/group. All data shown are mean ± s.e.m

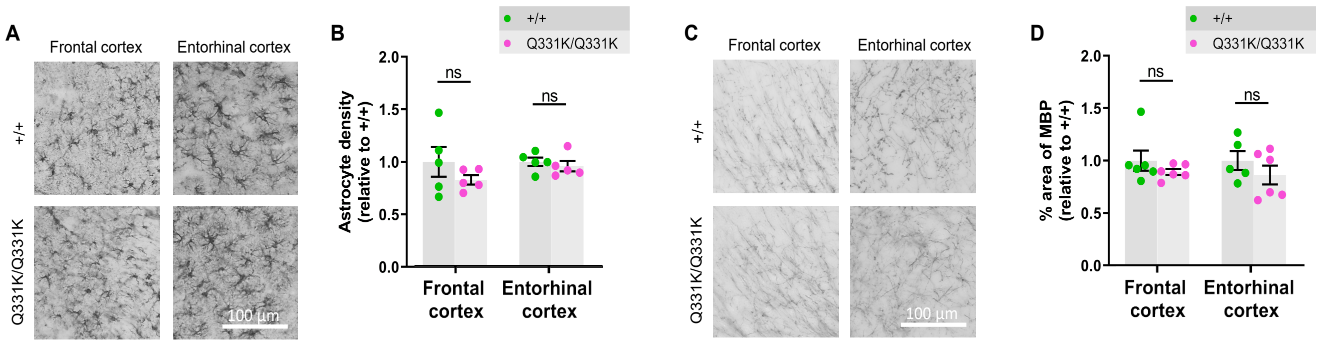


**Supplementary fig. 3 No significant changes in astrocyte density and MBP % area in TDP-43$^{Q331K}$ mice. A** Representative images showing astrocyte staining (dark brown) in given regions of 7-month-old +/+ and Q331K/Q331K mice. **B** Quantification of astrocyte density based on GFAP immunoreactivity comparing +/+ to Q331K/Q331K mice: frontal cortex, ns *P*= 0.4801; and entorhinal cortex, ns *P*=0.5450. **C** Representative images showing MBP staining (dark brown) in given regions of 7-month-old +/+ and Q331K/Q331K mice. **D** Quantification of % area of MBP comparing +/+ to Q331K/Q331K: frontal cortex, ns *P*= 0.5190; entorhinal cortex, ns *P*=0.5190. (**B, D**)*P* values were calculated with multiple t-tests adjusted by Holm-Sidak correction. n=5-6/group. All data shown are mean ± s.e.m

**Supplementary Table 1. Antibodies used in immunohistochemistry.**

| Protein | Primary Ab (product No.) | Primary Ab (dilution) | Secondary Ab (product No.) | Secondary Ab (dilution) |
|---|---|---|---|---|
| PV | Mouse, Sigma P3088 | 1:200 | Goat, Alexa Fluor 488, Thermo Fisher | 1:500 |
| | Rabbit, Abcam ab11427 | 1:1000 | Goat, Alexa Fluor 488, Thermo Fisher | 1:500 |
| GAD67 | Mouse, Chemicon MAB5406 | 1:1000 | Donkey, Alexa 568, Thermo Fisher | 1:500 |
| Iba1 | Goat, Abcam ab5076 | 1:500 | Donkey, Alexa Fluor 488, Thermo Fisher | 1:500 |
| Tmem119 | Rabbit, Abcam ab209064 | 1:1000 | Goat, Alexa Fluor 568, Thermo Fisher | 1:500 |
| Ki67 | Rabbit, Abcam ab16667 | 1:500 | Goat, biotinylated, Vector Lab BA1000 | 1:400 |
| DCX | Goat, Santa Cruz Biotechnology | 1:200 | Horse, biotinylated, Vector Lab BA9500 | 1:400 |
| SOM | Rat, Chemicon YC7 | 1:100 | Goat, biotinylated, Vector Lab BA9401 | 1:500 |
| MBP | Rat, Abcam ab7349 | 1:1000 | Goat, biotinylated, Vector Lab BA9401 | 1:500 |
| GFAP | Rabbit, Dako Z0334 | 1:2000 | Goat, biotinylated, Vector Lab BA1000 | 1:1000 |

**Supplementary table 2 MRC numbers and conditions of patients.**

| ALS patients | | | | |
|---|---|---|---|---|
| **MRC numbers** | **PMI (hours)** | **Age (years)** | **Gender** | **Genetics** |
| BNN_20604 | 26 | 62 | F | C9ORF72+ve |
| BNN_20993 | 55 | 43 | M | C9ORF72+ve |
| BNN_20613 | 89 | 50 | M | C9ORF72+ve |
| BNN_30081 | 63 | 73 | M | Negative |
| BNN001.28407 | 93 | 67 | F | Negative |
| BNN001.30220 | 70 | 56 | M | Negative |
| BNN_29694 | 102 | 50 | M | Negative |
| BNN001.26126 | 119 | 71 | M | Negative |
| BNN001.26497 | 84 | 63 | F | C9ORF72+ve |
| BNN001.28409 | 93 | 62 | F | C9ORF72+ve |
| BNN001.28792 | 51 | 68 | M | C9ORF72+ve |
| BNN_30175 | 84 | 69 | F | Negative |
| BNN001.28790 | 105 | 70 | M | Negative |
| BNN001.26765 | 30 | 66 | F | Negative |
| BNN_22225 | 109 | 62 | M | Negative |
| BNN_18803 | 96 | 75 | F | Negative |
| BNN001.26729 | 131 | 83 | F | Negative |
| BNN001.31439 | 59 | 62 | M | Negative |
| Means | 81 | 64 | N/A | N/A |
| PMI = post mortem interval; MRC = Medical Research Council. | | | | |

**Supplementary table 3 MRC numbers and conditions of controls.**

| Neurologically healthy individuals | | | |
|---|---|---|---|
| **MRC numbers** | **PMI (hours)** | **Age (years)** | **Gender** |
| BNN_19686 | 75 | 77 | F |
| BNN001.28960 | 126 | 37 | F |
| BNN001.28402 | 49 | 79 | M |
| BNN001.28495 | 39 | 78 | M |
| BNN001.28959 | 86 | 39 | M |
| BNN001.29525 | 99 | 52 | M |
| BNN001.28794 | 72 | 79 | F |
| BNN001.29526 | 67 | 47 | M |
| BNN001.28797 | 57 | 79 | M |
| BNN001.28406 | 72 | 79 | M |
| BNN001.28793 | 72 | 79 | F |
| Means | 74 | 66 | N/A |
| PMI = post mortem interval; MRC = Medical Research Council. | | | |
